# DNA methylome-based validation of induced sputum as an effective protocol to study lung immunity: construction of a classifier of pulmonary cell types

**DOI:** 10.1101/2021.03.12.435086

**Authors:** Jyotirmoy Das, Nina Idh, Liv Ingunn Bjoner Sikkeland, Jakob Paues, Maria Lerm

**Author notes:** Emails: Jyotirmoy Das, Nina Idh, Jacob Paues, Liv Ingunn Bjoner Sikkeland, Maria Lerm. Shared first authorship (Jyotirmoy Das, Nina Idh). Corresponding author:* Maria Lerm, Professor in Medical Microbiology, Div. of Infection and Inflammation, Lab 1, floor 12, Dept. of Biomedical and Clinical Sciences, Faculty of Medicine and Health Sciences, Linköping University, SE-58185 Linköping, Sweden.

## Abstract

**Background:** Flow cytometry is a classical approach used to define cell types in peripheral blood. While DNA methylation signatures have been extensively employed in recent years as an alternative to flow cytometry to define cell populations in peripheral blood, this approach has not been tested in lung-derived samples. Here, we compared bronchoalveolar lavage with a more cost-effective and less invasive technique based on sputum induction and developed a DNA methylome-based algorithm that can be used to deconvolute the cell types in such samples.

**Results:** We analyzed the DNA methylome profiles of alveolar macrophages and lymphocytes cells isolated from the pulmonary compartment. The cells were isolated using two different methods, sputum induction and bronchoalveolar lavage. A strong positive correlation between the DNA methylome profiles of cells obtained with the two isolation methods was observed, and in two of the donors, in which the correlation was best, a later analyses demonstrated that those subjects the samples were consistently derived from the lower part of the lungs. We also identified unique patterns of CpG methylation in DNA obtained from the two cell populations, which can be used as a signature to discriminate between the alveolar macrophages and lymphocytes by means of open-source algorithms. We validated our findings with external data and obtained results consistent with the previous findings.

**Conclusions:** Our analysis opens up a new possibility to identify different cell populations from lung samples and promotes sputum induction as a tool to study immune cell populations from the lung.

## Background

Much of our knowledge of the immune system is derived from studies on cells isolated from blood, although extrapolation of the performance of those lymphocytes and myeloid cells gives very limited information on immunological events in peripheral tissues. It is becoming increasingly evident that in order to understand tissue-specific immunity, samples from relevant tissues have to be collected and studied. To understand how different environmental factors contribute to airway inflammation, sputum induction (SI) has been employed as replacement for the invasive bronchoalveolar lavage (BAL) in several studies [1–4].

In this report, we isolated both macrophages (HLA-DR+, CD3-cells) and lymphocytes (CD3+ cells) from samples obtained through SI and BAL from the same donors in order to compare the cell populations obtained through the two protocols. Instead of using the marker-biased flow cytometry approach for cell characterization, we employed whole genome DNA methylome analyses for the characterization. Our data demonstrate that the DNA methylation profile of the cell populations display strong overlap and conclude that the SI approach represents a valuable tool for non-invasive collection of samples representing the mucosal immunity of the airways. In addition, we provide a classifier algorithm that, based on DNA methylome data, can distinguish macrophages in samples derived from the pulmonary compartment.

## Results

### DNA methylomes from cells isolated through SI and BAL are highly correlated

To investigate whether the pulmonary cells collected using SI are similar to those obtained through BAL, we collected both specimens from the same subjects and isolated lymphocytes (CD3+) and macrophages (HLA-DR+/ CD3-). DNA methylomes were captured from the isolated DNA using the Illumina 450K protocol (**Figure 1**). The variation (β values after the data filtration) in the DNA methylome datasets derived from cells from SI and BAL were calculated and a Spearman’s rank correlation test from each cell populations and each subject were preformed.. The analysis revealed a strong and highly significant positive correlation (*r*IS|BAL x□=0.95, *p* value < 2.2e-16) in each pair of data obtained through the two isolation protocols (**Figure 2, Table 2, Supplementary Figure S1a**), suggesting that the cell identities and phenotypes were very similar. To analyze any possible common differences between sputum cells and BAL cells, we compared the mean global β values for the respective cell type and found that in both cell populations, the values differed slightly and significantly between the two protocols (SI_mean_ = 0.4504, BAL_mean_ = 0.4669 in HLA-DR+/CD3− cells and SI_mean_ = 0.4727, BAL_mean_ = 0.4666 in CD3+ cells, **Figure 3a and b**), suggesting that there is a relevant difference between the two protocols. Of note, a previous study demonstrated that the DNA methylomes of macrophages recovered from lower parts or apical parts of the lung display differences in the DNA methylomes, probably reflecting differences in the phenotypes [5]. Therefore, to investigate whether our HLA-DR data could be used to predict from which part of the lung the SI-derived cell populations came from, we used previously published data from Armstrong *et al* (GSE132547) that consists of Illumina EPIC array 850K data obtained from 12 healthy subjects and grouped them separately as upper and lower lung for each subject. The linear regression was calculated and residuals were estimated 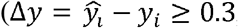, the distance of the β value of each CpG from the linear regression line, **Supplementary Figure S1b**) to obtain the CpGs which are at the farthest distance from the regression line and strongly correlated to either SI or BAL samples. The percentage of cells derived from the upper and/or lower lungs was identified by using extracted CpGs from the HLA-DR datasets as test data while using the Armstrong *et al* data [5] as training data sets in the EpiDISH package. For the subjects P4 and P5, the the analysis predicted that BAL recovered cells were from lower lung and SI cells were from the upper lung, wheras for P2 and P3, SI and BAL cells were identified as cells from the lower lung. In P1, a fraction of the BAL cells (21%) were identified from the upper lung while the rest of the BAL cells and the SI cells from the lower lung.

**Figure 1:**
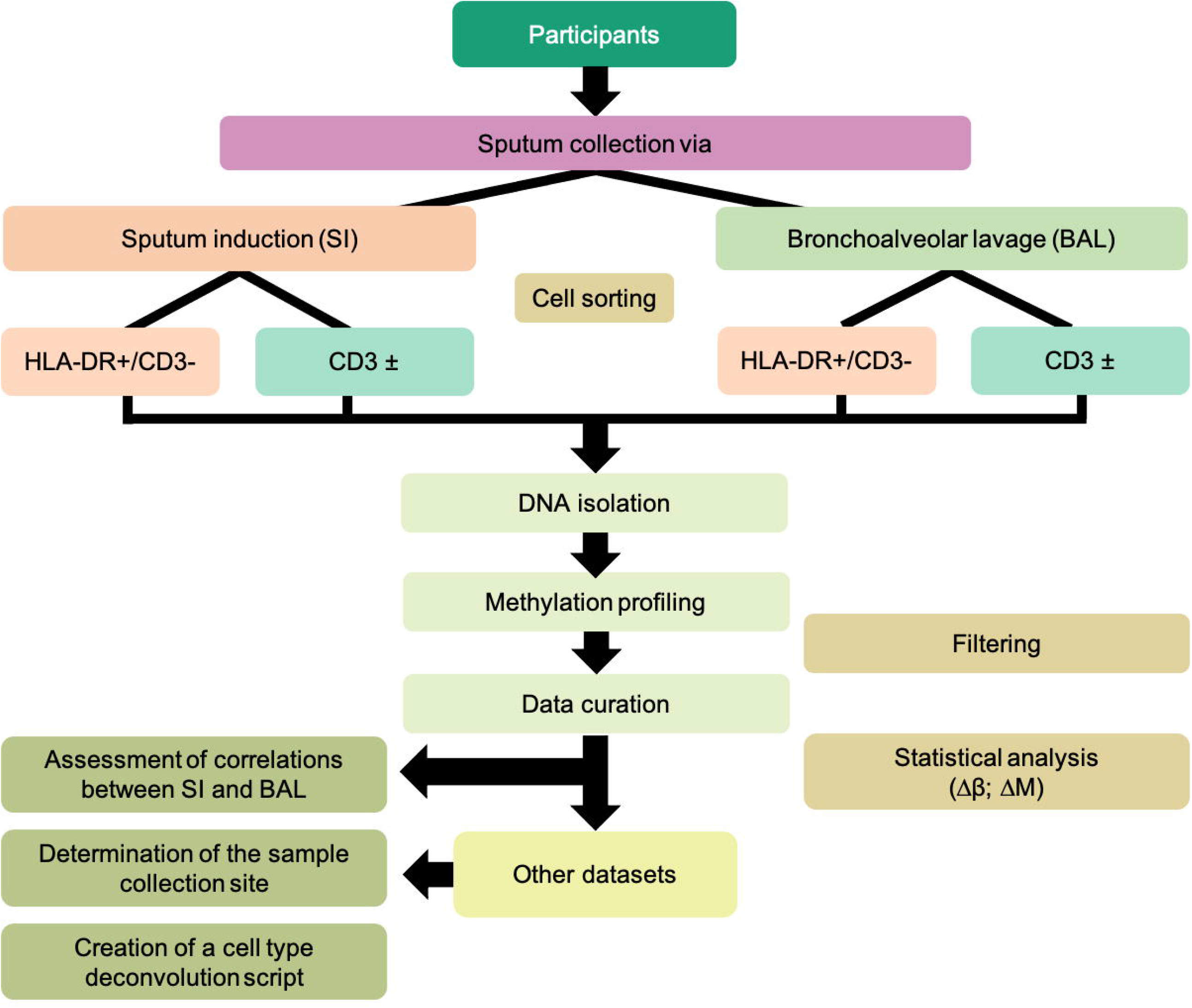
Flow diagram of the study.

**Figure 2:**
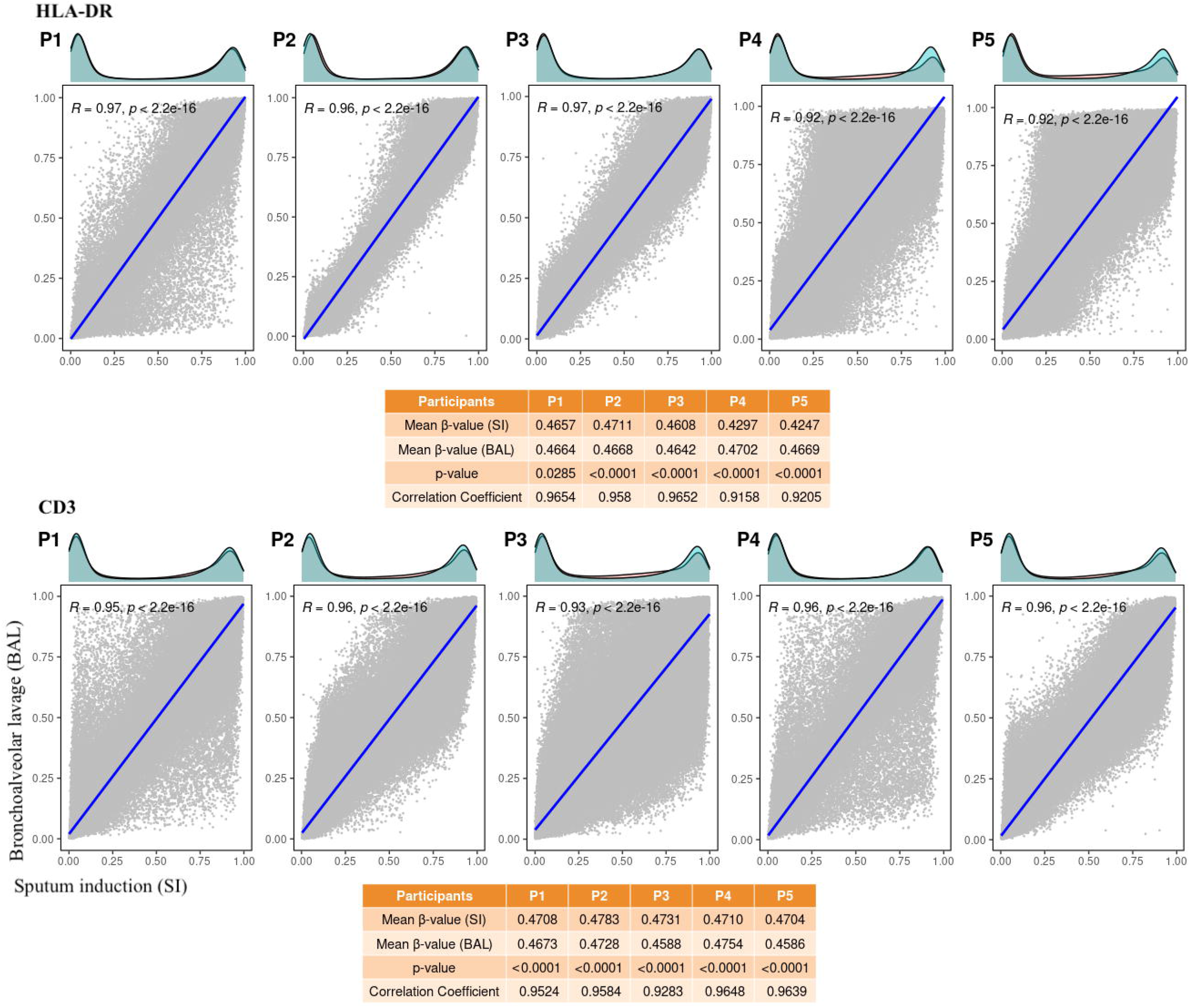
Spearman’s rank correlation analysis between sputum induction (SI) and bronchoalveolar lavage (BAL) in HLA-DR and CD3 cell population for each patient. The blue line is the linear regression line. Each dot illustrates one CpG. The axes represent SI (x-axis) and BAL (y-axis). The top layer of the correlation plot depicts the density plot to display the similarity/dissimilarity among SI and BAL methods. The adjacent table represents statistical data from each sample. The upper plot represents data from HLA-DR cells and the lower plot shows the data analysis result from CD3 cells.

**Figure 3:**
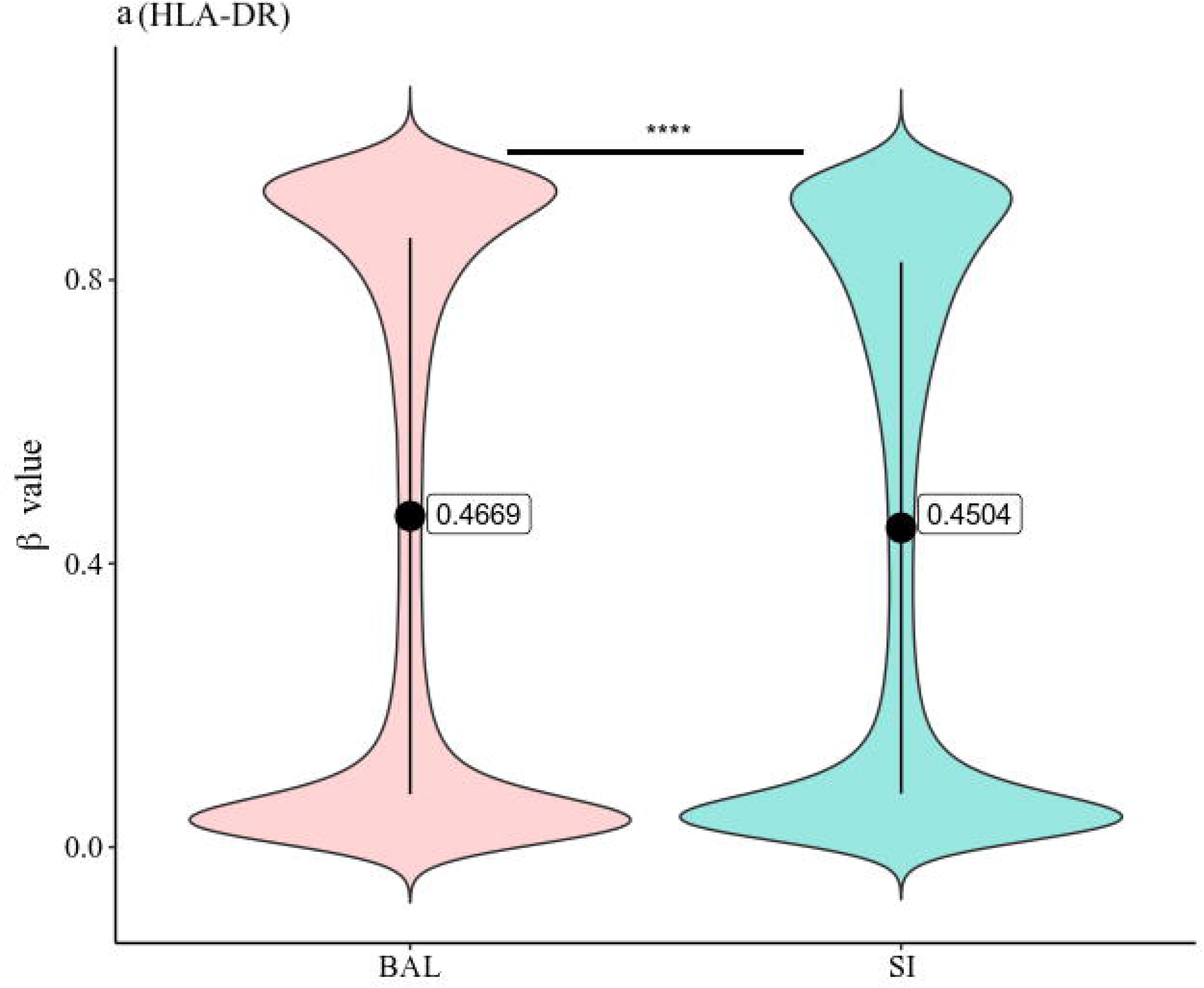
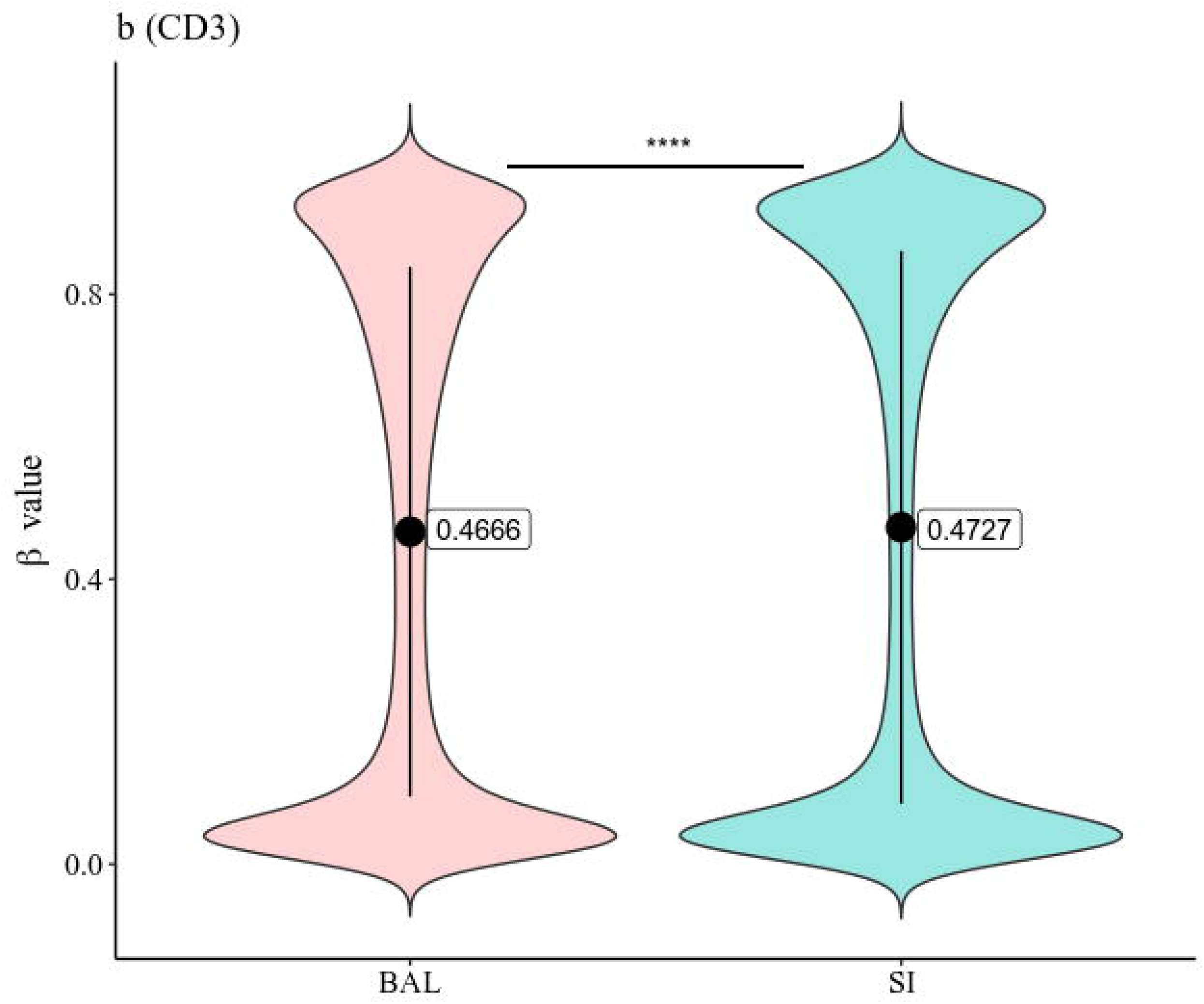
Distribution of DNA methylation β value in SI and BAL data (HLA-DR and CD3). The y-axis of bean plot represents the distribution of β values of SI and BAL. The mean value is indicated using the black solid circle and the vertical line represents the range of the β value. a. represents the β value distribution from HLA-DR cells and b. represents the β value distribution from CD3 cells. Statistics: Mann-Whitney-Wilcoxon test, **** represents a *p*-value <1e-4. N_HLA-DR_ = 382,591, N_CD3_ = 398,390.

**Figure 4:**
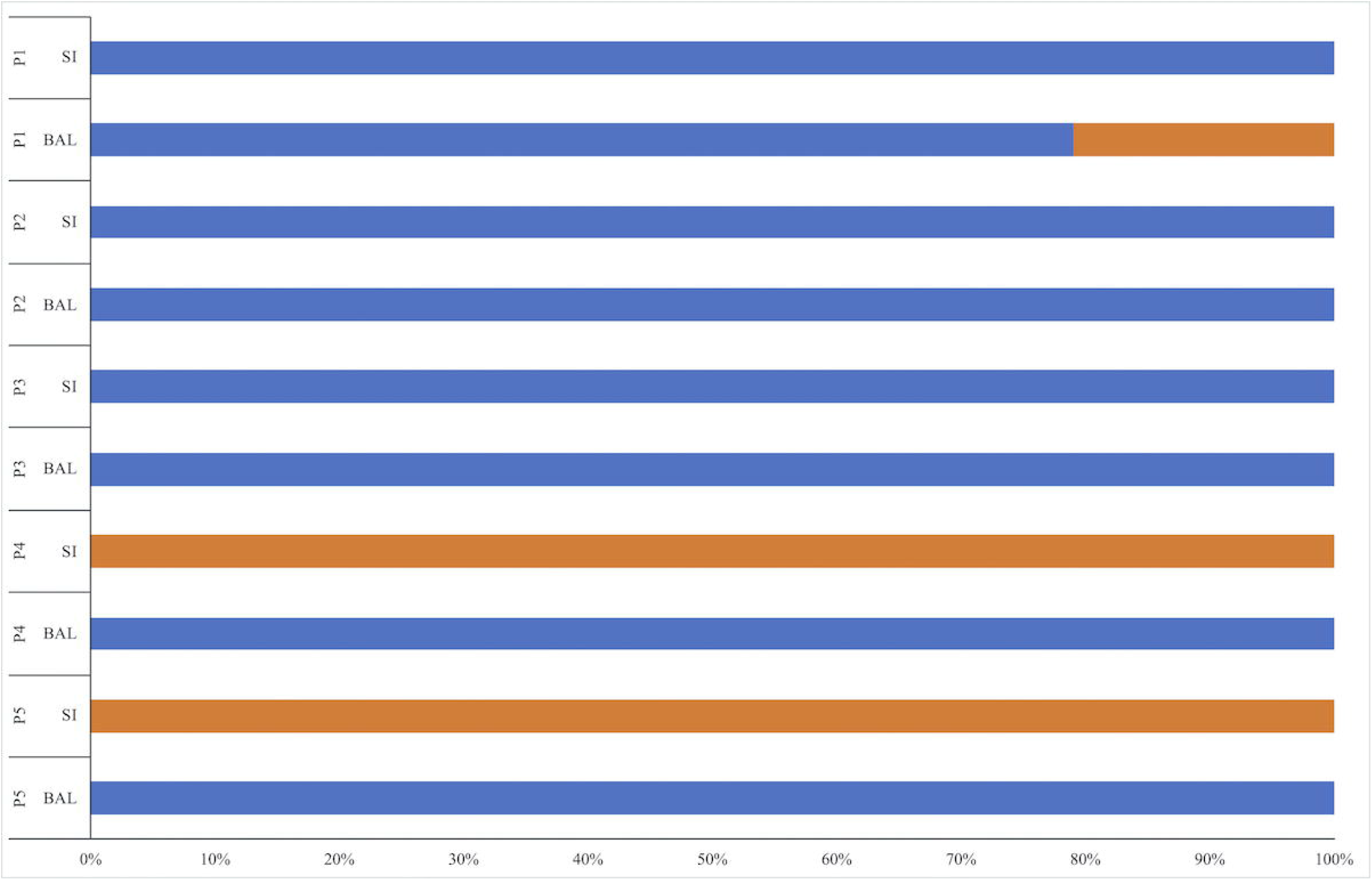
The horizontal stacked bar plot displays the distribution of CpGs in lower and upper right lungs in each sample for two different isolation methods. Blue and orange color represents lower and upper lungs, repectively. The x-axis shows the percentage of CpGs present in the respective area of the right lung.

### A DNA methylome-based cell-sorting algorithm for pulmonary samples

In 2012, Houseman *et al* presented a cell-sorting algorithm based on DNA methylation data to identify the fraction of five different cell types in peripheral blood mononuclear cells (PBMCs) [6]. The classifier has had a fundamental impact on deconvolution of blood-derived DNA methylome data (nearly 2000 citations as per March 2021, including [7–12]) and has also been extrapolated to pulmonary cell DNA methylomes [13]. The datasets compiled in our present study constitute a relevant source of data to create a similar classifier tailored for the pulmonary compartment.

Based on our demonstration that the DNA methylomes from both isolation procedures (SI and BAL) were highly similar, we averaged the SI and BAL data for each cell population (HLA-DR and CD3) as a first step. To identify cell type-specific CpGs, we employed three different reference-free EWAS algorithms, *RefFreeEWAS* [14], *SVA* [15] and *RefFreeCellMix* [14]. To estimate the variation only from the strongly correlated CpGs, we again set the residuals, 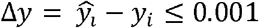, and identify those CpGs close to the regression line to avoid SI or BAL-specific CpGs (**Supplementary Figure S1b**). Using the above-mentioned algorithms (with *p*_BH_-value < 0.05), we determined the CpGs specific for HLA-DR+ and CD3+ cells and (**Table 5**, see **methods**) identified a total of 594 CpGs and 2,292 CpGs for HLA-DR cells and CD3 cells, respectively (**Figure 5a, b**). Finally, after applying an additional cut-off (≤ 0.1) to remove CpG with only minor differences in β values, we compiled a total of 2,763 CpGs as the reference dataset for the alveolar cell sorting (**Supplementary Table S1**).

**Figure 5:**
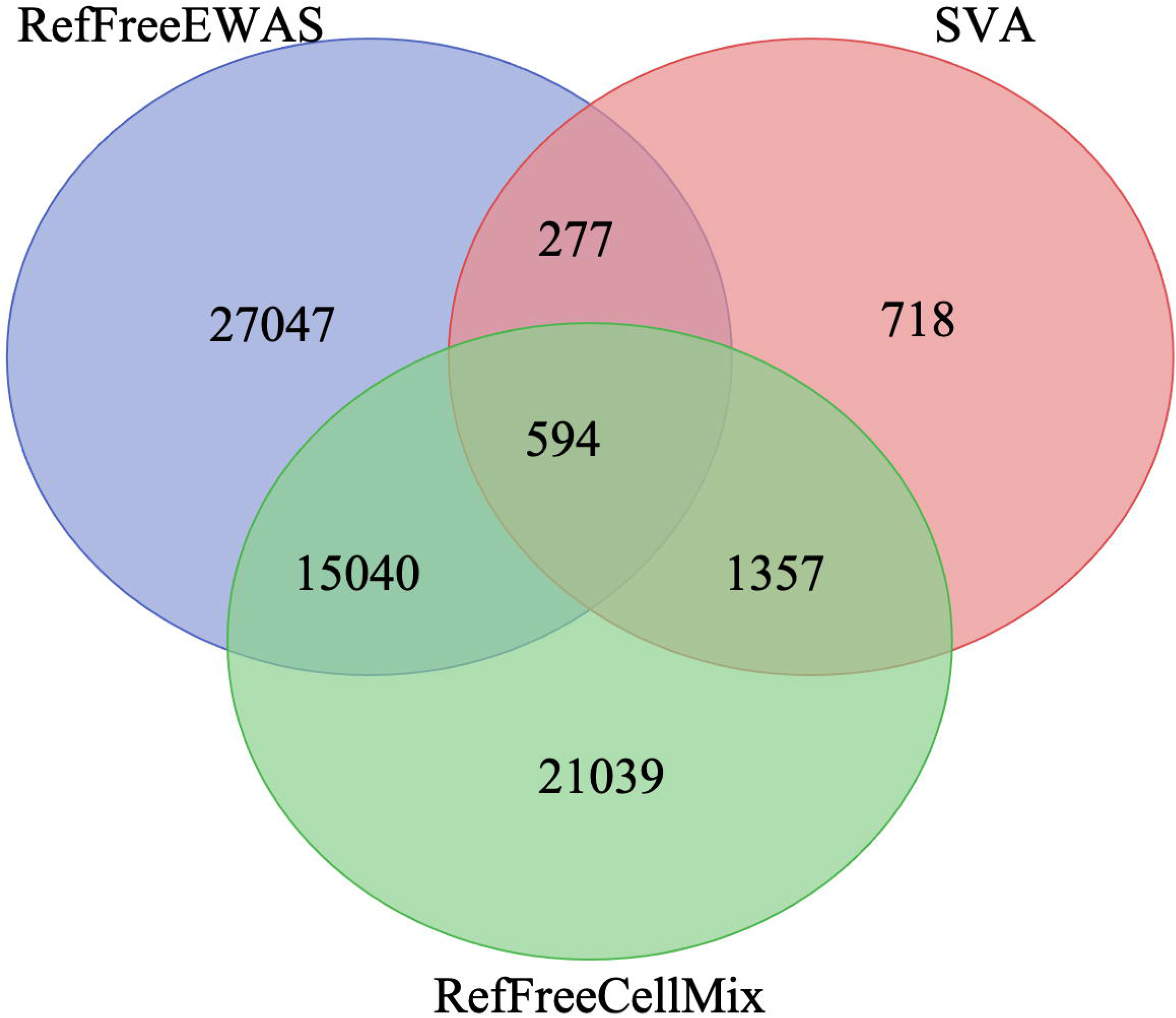
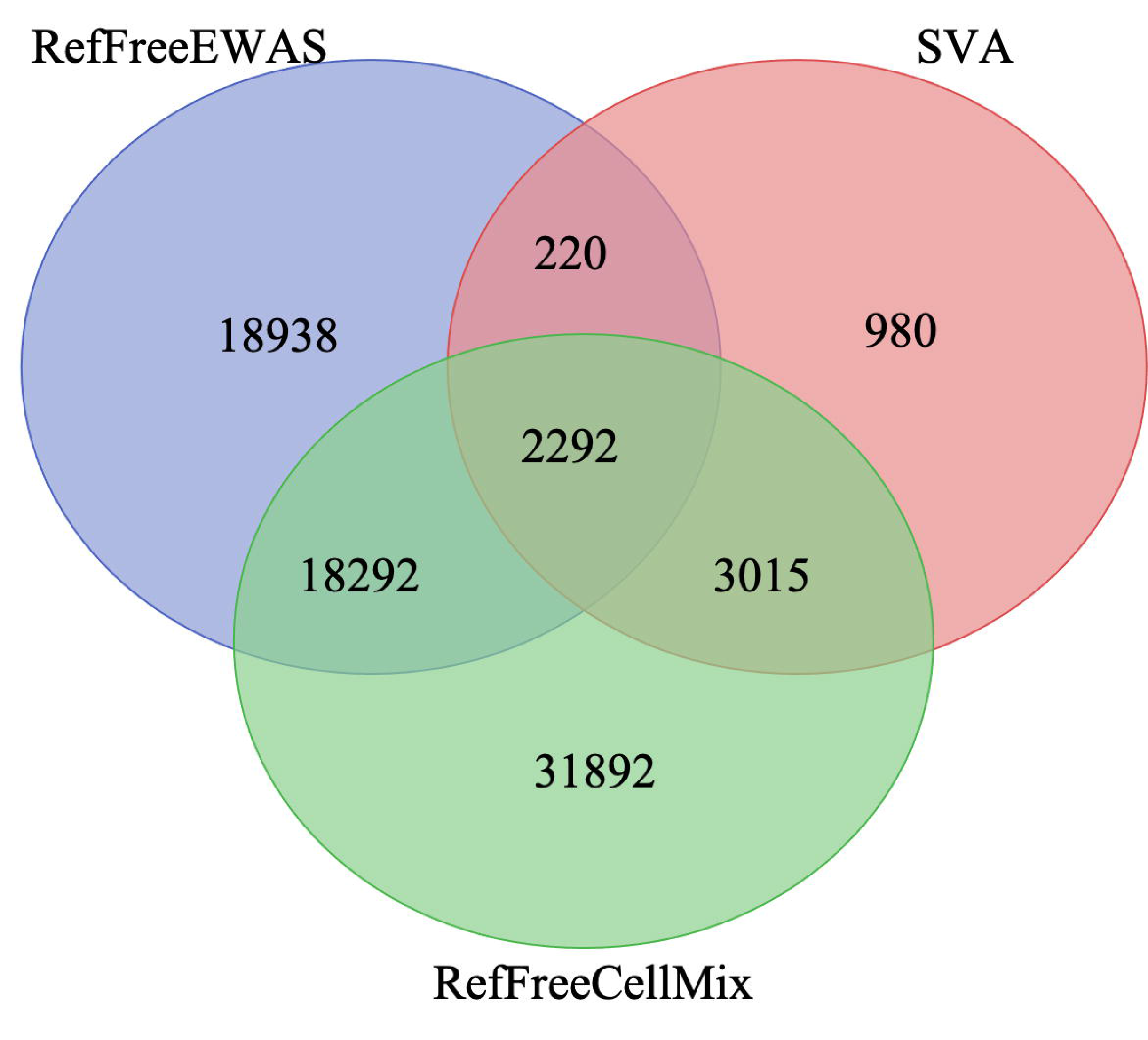
The Venn diagrams in **a** and **b** represent HLA-DR specific common CpGs and CD3 specific common CpGs based on three different reference free EWAS deconvolution methods, RefFreeEWAS, SVA and RefFreeCellMix, respectively.

### Validation of the classifier against another dataset reveals a high accuracy

In order to validate the identified cell-specific CpGs, we compared the data with a recently published BAL sample DNA methylome dataset (GSE133062) by Ringh *et al* [13]. In this study, the authors demonstrated by analyzing Giemsa-stained BAL cytospins that more than 90% of the cells were alveolar macrophages. We performed a cell proportion analysis using our classifying CpG set in combination with the *EpiDISH* package in R with the whole 850K DNA methylation data of Ringh *et al* [13], The results revealed a high accuracy of the reference data in all samples with a high proportion of alveolar macrophages (85-100%) and low proportion of alveolar lymphocytes (0-15%) (**Figure 6**) in line with the published data [13], The computational validation of the result showed that the identified CpGs can be a potential signature to deconvolute cell proportions in lung samples based on DNA methylation data.

**Figure 6:**
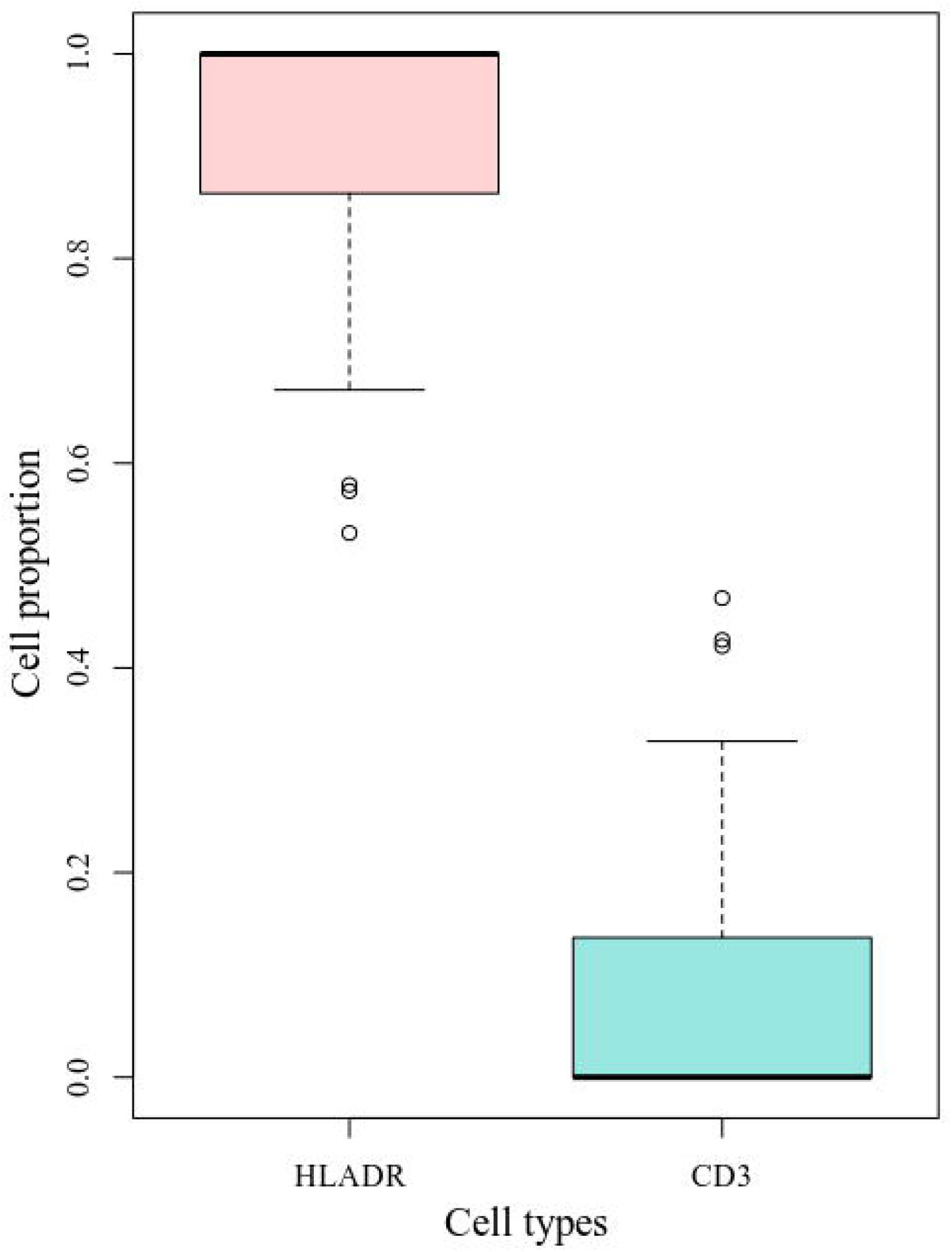
Boxplot illustrates the use of cell sorting algorithm to analyse the cell proportion from the cell specific CpGs identified from the BAL samples of the current study and compared with the Ringh *et al* (2019) 850K data (GSE133062). The pink colour showed the alveolar macrophages and sea-green colour for the alveolar lymphocytes. The y-axis scale denotes the proportion of cells, ranging from 0 to 1.

## Discussion

In this study, we demonstrated a strong correlation between the DNA methylome profiles of two cell populations isolated from SI and BAL. Both isolation methods are well known and have been used for decades for clinical diagnosis of pulmonary conditions. The sensitiveness of both methods has been widely compared [16-18] and one obvious advantage with BAL is the option to collect samples from defined sites in the lung. However, a major disadvantage with BAL is that it is an invasive procedure performed by physicians at larger medical centers that requires sedation and monitoring of the patient. It is also considerably more expensive than SI. On the other hand SI is a safe, relatively cheap, non-invasive technique that can be performed by nurses or physiotherapists at an out-patient clinic, which all contributes to the limitation of BAL samples for research [19].

To explore the comparativeness of cells obtained through SI and BAL, we collected samples from five patients using both methods and isolated HLA-DR+/CD3- and CD3+ cells. We performed a genome-wide DNA methylation analysis to compare the β values from ~400k CpG sites in the DNA from these cell types and found a strong and significant correlation between these two methods in both cell types. Although this analysis suggested that the cell populations obtained through the two protocols were very similar, comparison of the global mean β values significantly differed between SI and BAL. Through exploration of a previous study’s finding [5] that designated unique CpG signatures to macrophages collected from the upper or lower parts of the lungs, we examined the possibility that this slight discrepancy was related to the local pulmonary site of sample collection. Indeed, the analysis allowed prediction of the site of collection (either upper or lower lung or in one case a mixture of both). Importantly, like BAL, which allows sample collection at defined sites, SI samples could result in cells obtained from the lower parts, which demonstrates that SI is an effective method for studies of pulmonary immunity.

Using DNA methylation-based signatures to classify cell types is gaining increasing interest. For example, a recent study showed that tissue-specific DNA methylation signatures of free circulating DNA in plasma of patients with severe Covid-19 can reveal specific organ injury [20]. The most widely used cell type deconvolution algorithm was described by Houseman *et al* [6], which uses cell type-specific DNA methylation signatures to determine the frequencies of T cells, B cells, monocytes, NK cells and neutrophils in PBMCs. Here, we present a comparable algorithm to assess the proportion of T cells and macrophages in pulmonary samples by combining three reference-free algorithms to derive the cell-specific CpGs. We used reference-free algorithms since there is no available reference data. We observed a large variation among these three algorithms and therefore intersected the CpG list and used the overlapping CpGs as cell type markers. In a validation step, our algorithm was applied to another publicly available dataset and it generated a similar prediction of cell proportions as the previously published, microscopy-validated data. The results has the potential to facilitate interpretation of DNA methylation data originating from pulmonary samples, which is receiving increasing attention with a large number of studies defining pulmonary CpG signatures for conditions including cystic fibrosis, lung cancer, chronic obstructive pulmonary diseases and smoking [13,21–24].

## Conclusions

Our analyses demonstrate that SI is an attractive method for studies of pulmonary immunity that can replace BAL and that DNA methylomes derived from the cells obtained through this protocol can distinguish between cells collected from the upper or lower parts of the lungs as well as predict the proportion of macrophages and T cells.

## Methods

### Study design, ethics and participants

Participants, five female patients scheduled for BAL as part of their clinical investigation, were recruited from Linköping University Hospital between 2017-2018 (**Table 1**). The participants donated induced sputum at a time point −/+ 2 weeks before/after collection of the BAL sample. After collection of BAL for diagnostic purposes, leftovers were used for the planned research.

**Table 1:** Demographic characteristics of participants

| Characteristics | Value |
| --- | --- |
| Age (year)<sup>†</sup> | 55±10 |
| Height (cm)<sup>†</sup> | 166±1 |
| Weight (kg)<sup>†</sup> | 70±10 |
| Body Mass Index (BMI)<sup>†</sup> | 25±3 |
| Sex (male/female) | 0/5 |
| BCG (yes/no) | 5/0 |
| Smoking (current/previous/never) | 0/4/1 |

<sup>†</sup> Values are mean ± standard deviation.

**Table 2:** Comparison between bronchoalveolar lavage (BAL) and sputum induction (SI)

| Participants | Spearman's rank correlation coefficient (*r*) |  | *p*-value |
| --- | --- | --- | --- |
|  | HLA-DR<br>(N = 382,591) | CD3<br>(N = 398,390) |  |
| P1 | 0.9654 | 0.9524 | < 2.2e-16 |
| P2 | 0.958 | 0.9584 |  |
| P3 | 0.9652 | 0.9283 |  |
| P4 | 0.9158 | 0.9648 |  |
| P5 | 0.9205 | 0.9639 |  |

The table describes the Spearman's rank correlation coefficient values from each participant in both HLA-DR and CD3 cell types. The observed *p*-value < 2.2e-16 for all data. The calculation was done in 382,591 CpGs in HLA-DR and 398,390 CpGs in CD3 cell types.

### Sputum induction and sputum processing

Sputum induction and processing was done as described by Sikkeland *et al* (2012) [25,26] with some modifications. Briefly, hypertonic saline at 3%, 4%, and 5% were inhaled for three 7-minutes periods using a nebulizer (eFlow Rapid, PARI, Germany) after nebulizer-administered premedication with an adrenergic β2-agonist, salbutamol 1ml (1mg/ml). After each inhalation period expectorates were collected, the subjects performed a three-step cleansing procedure to reduce contamination from nasopharynx (blow nose, rinse mouth and gargle three times with water), before coughing deeply involving the thorax and spitting the expectorate into a 50 ml sterile tube, (Falcon).

Sputum was processed within two hours after induction. From the sputum sample, mucus plugs, were manually selected, weighed and treated with four times its volume of 0.1% dithiothreitol (DTT, Thermo Fisher). The sample was aspirated with a Pasteur pipet five times, vortexed 15 sec and rocked at 4°C at 40 rev/min for 15 min. The sample was then diluted four times with phosphate buffered saline (PBS), and rocked for 5 min before filtrated through a 50 μm cell strainer (CellTrics^®^ by Sysmex) and centrifuged for 5min. Cell viability, total cell counts, and squamous cell counts were manually evaluated by counting cells in a Bürker chamber with trypan blue staining.

BAL was obtained according to standard clinical procedures at the Linköping University Hospital. 125 ml of sterile, physiological sodium chloride (NaCl) solution was instilled and aspirated to collect cells. The BAL fluid was kept on ice until processing. PBS was added and the samples were centrifuged at 400g for 5 minutes at 4°C, then washed with PBS and followed by recentrifuged at 300g for 13 minutes at 4°C before isolation of DNA as described below.

### DNA extraction and Data processing

The pulmonary HLA-DR and CD3-positive cells were isolated using superparamagnetic beads coupled with anti-human CD3 and Pan Mouse IgG antibodies (Invitrogen Dynabeads^®^, Life Technologies AS, Norway) and HLA-DR/human MHC class II antibodies (Invitrogen Dynabeads^®^, Life Technologies AS, Norway). An initial positive selection was done with CD3 beads followed by a positive HLA-DR selection. Bead coating and cell isolation was performed according to manufacturer’s protocol. The DNA and RNA were extracted from the lung immune cells using the AllPrep^®^ DNA/RNA Mini Kit (Qiagen, Germany), per the manufacturer’s instructions.

The DNA methylome data of HLA-DR+/CD3- and CD3+ cells was analyzed using the HumanMethylation450K (450K) BeadChip (Illumina, USA) array as per the manufacturer’s instructions. The raw IDAT files of the DNA methylation data was processed using the BMIQ normalization function of the *ChAMP* package [27] in R (v3.6.3) after using a robust filtration criteria to generate the β values from each CpG for each sample. The sex chromosome data was removed to get rid of the gender biasness, multi-hit probes were taken away from the data and also CpGs with lower *p*-values (< 0.01) was drawn out from the data ([28]; submitted manuscript). The filtered data was normalized using the beta-mixture quantile normalization (BMIQ) function using the ChAMP package [29]. To compare our 450K data with other previously published dataset from Illumina 850K array [GSE133062, GSE132547 [5,13]], we used *merge* function in R to extract the CpG sites present in the Illumina 450K array analysis.

### Statistical calculations

All statistical analysis was performed in R v3.6.3 and bioconductor packages (v3.10). Anderson-Darling (AD) normality test [30] was used to calculate the non-parametric distribution of the dataset using *nortest* package [31] in R (used for large dataset, >380K, **Table 3**). As all of the data show non-parametric distribution, Spearman’s rank correlation test was performed to calculate the correlation using *ggpubr* package [32] in R. The linear regression line with *geom smooth* function from *ggplot2* package was added to the plot to calculate the confidence interval (C.I. = 95%). The correlation matrix was calculated using the Pearson’s correlation coefficient to evaluate the pairwise relationship among all samples. The value was considered significance if *p*-value < 0.05 throughout the current study.

**Table 3:**
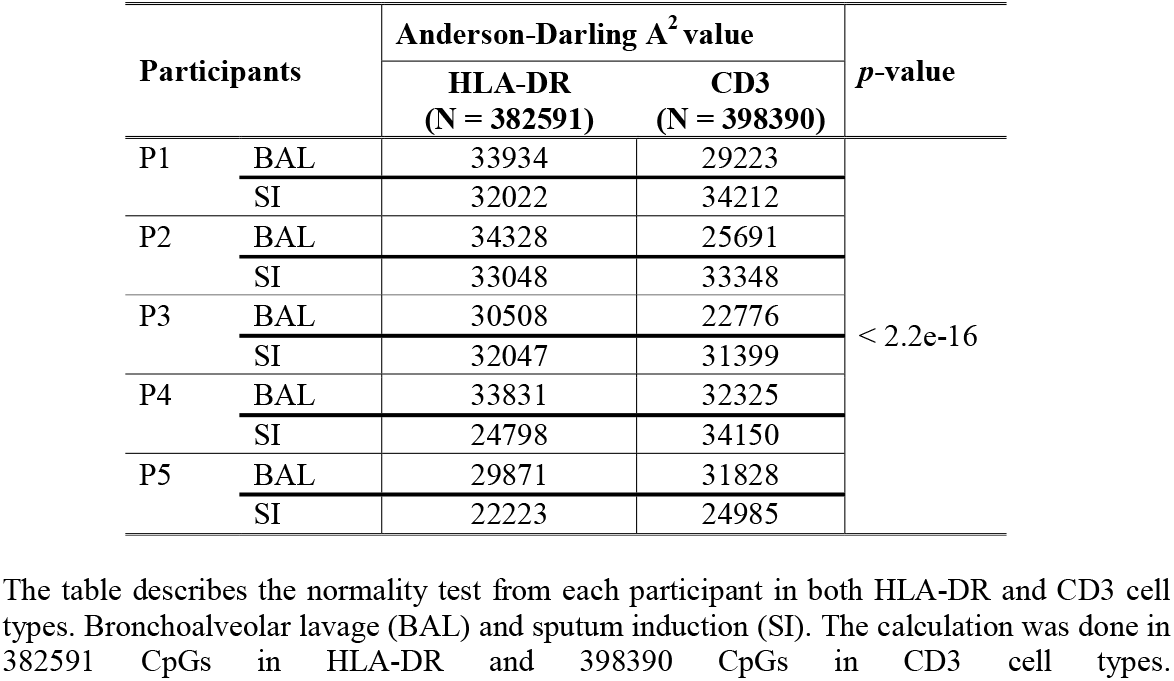
Normality test result from Anderson-Darling test

**Table 4:** Independent sample test Mann-Whitney-Wilcoxon rank sum test

| Participants | Mann-Whitney-Wilcoxon rank sum test |  |  |  |  |  |
| --- | --- | --- | --- | --- | --- | --- |
|  | HLA-DR (N= 382,590) |  |  | CD3 (N = 398,389) |  |  |
| | SI<br>(mean $\beta$) | BAL<br>(mean $\beta$) | *p*-value | SI<br>(mean $\beta$) | BAL<br>(mean $\beta$) | *p*-value |
| P1 | 0.4657 | 0.4664 | 0.0285 | 0.4708 | 0.4673 | 1.168e-09 |
| P2 | 0.4711 | 0.4668 | < 2.2e-16 | 0.4783 | 0.4728 | < 2.2e-16 |
| P3 | 0.4608 | 0.4642 | < 2.2e-16 | 0.4731 | 0.4588 | 4.518e-06 |
| P4 | 0.4297 | 0.4702 | < 2.2e-16 | 0.4710 | 0.4754 | < 2.2e-16 |
| P5 | 0.4247 | 0.4669 | < 2.2e-16 | 0.4704 | 0.4586 | < 2.2e-16 |

Bronchoalveolar lavage (BAL) and sputum induction (SI). The calculation was done in 382591 CpGs in HLA-DR and 398390 CpGs in CD3 cell types.

### Cell-specific CpGs identification and validation

To identify the cell type-specific CpGs, three reference-free algorithms were used, *RefFreeEWAS* [14], *SVA* [15] and *RefFreeCellMix* [14]. First, the linear regression model was applied on the β values for each sample combining both SI and BAL samples. The residual value was then calculated using the equation, 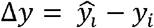, and Δ*y* < 0.001 filter was set to extract the CpGs close to the regression line (**Supplementary Figure S1b**). Data was prepared from the β values and the phenotypes (e.g., HLA-DR and CD3) per sample as per algorithm requirement and covariates were calculated using Bonferroni-Hochberg corrected *p*-value < 0.05. Each algorithm was used separately to predict the cell-type-specific CpGs (**Table 5**). Venn analysis was used to intersect the overlap CpGs identified using the three algorithms. To validate our result with another external dataset, a previously published dataset was used as the test data GSE133062 [13] only in the HLA-DR cell types using the *EpiDISH* package [33] in R. No publicly available dataset was identified that could be used to validate the CD3-specific cell population.

**Table 5:** Cell specific CpGs to determine cell proportion estimation using different algorithms

| **Algorithms** | **HLA-DR specific CpGs** | **CD3 specific CpGs** |
| --- | --- | --- |
| *RefFreeEWAS* | 42,958 | 39,742 |
| *SVA* | 2,946 | 6,507 |
| *RefFreeCellMix* | 38,030 | 55,491 |

RefFreeEWAS: Epigenome-Wide Association Study (EWAS) using Reference-Free DNA Methylation Mixture Deconvolution. SVA: Surrogate Variable Analysis and RefFreeCellMix: Reference-Free Cell Mixture Projection

## Declarations

### Ethics approval and consent to participate

The study was approved by the ethics committee, EPN, in Linköping, Dnr 2016/237-31. Oral and written informed consent were obtained from all participants before inclusion.

### Consent for publication

All authors have approved to the final version of the manuscript.

### Availability of data and materials

Illumina BeadChip 450K array data will be made available after final publication and the R scripts are available in github.com/JD2112/SIvsBALanalysis (upon request). The list of cell-type-specific CpGs are added in the **Supplementary Table S1**.

### Authors’ contributions

M.L. N.I. and J.P. wrote the ethical application, M.L., N.I., J.P. and J.D. conceived and designed the study, N.I. and J.P. collected the patient samples and compiled the demographics tables, L.I.B.S. designed the sputum induction method, J.D. and M.L. designed and performed the bioinformatic analysis of the data, J.D. wrote the scripts and created the figures, M.L., N.I. and J.D. wrote the manuscript.

## Acknowledgements

We direct our gratitude to the staff at Linköping University Hospital for assistance in sample collection and all the subjects for donating samples. The DNA methylome data were generated at the Bioinformatics and Expression Analysis (BEA) Core Facility at the Department of Biosciences and Nutrition, which is supported by the Board of Research at the Karolinska Institute, Stockholm. The computations were enabled by resources provided by the Swedish National Infrastructure for Computing (SNIC) at Linköping University campus partially funded by the Swedish Research Council through grant agreement no. 2018-05973.

---

**Supplementary Figure S1a:** Bean plots to validate the SI samples in compare to the BAL samples in both macrophages and lymphocytes cell types for cell specific CpGs. Pink colour shows the SI and sea-green colour for the BAL samples. The y-axis scale denotes the β value of the CpGs in samples. The x-axis shows the distribution in different sample. The horizontal black line at the middle represents the mean β value for each isolation method in each sample. a. represents the HLA-DR cells and b. show the data from the CD3 cells.

**Supplementary Figure S1b:** Figure representing the calculation of residual value. The tangent distance was calculated using the equation mentioned in **methods**. The ellipses were drawn to show the CpGs closer to SI and BAL samples. The dashed line represents the cut-off applied to extract the CpGs above Δ*y* = 0.3. Each dot reprensts one CpG.

**Supplementary Figure S2**: The correlation matrix among the SI and BAL samples in both alveolar macrophages and lymphocytes cells calculated using the cell specific CpGs. The numbers showed the pairwise pearson’s correlation coefficient. The colour scale shows the correlation coefficient values, ranging from −1 to 1.

