## Supplementary figures and images for "DNA methylome-based validation of induced sputum as an effective protocol to study lung immunity: construction of a classifier of pulmonary cell types"

### Supplementary Figure S1a

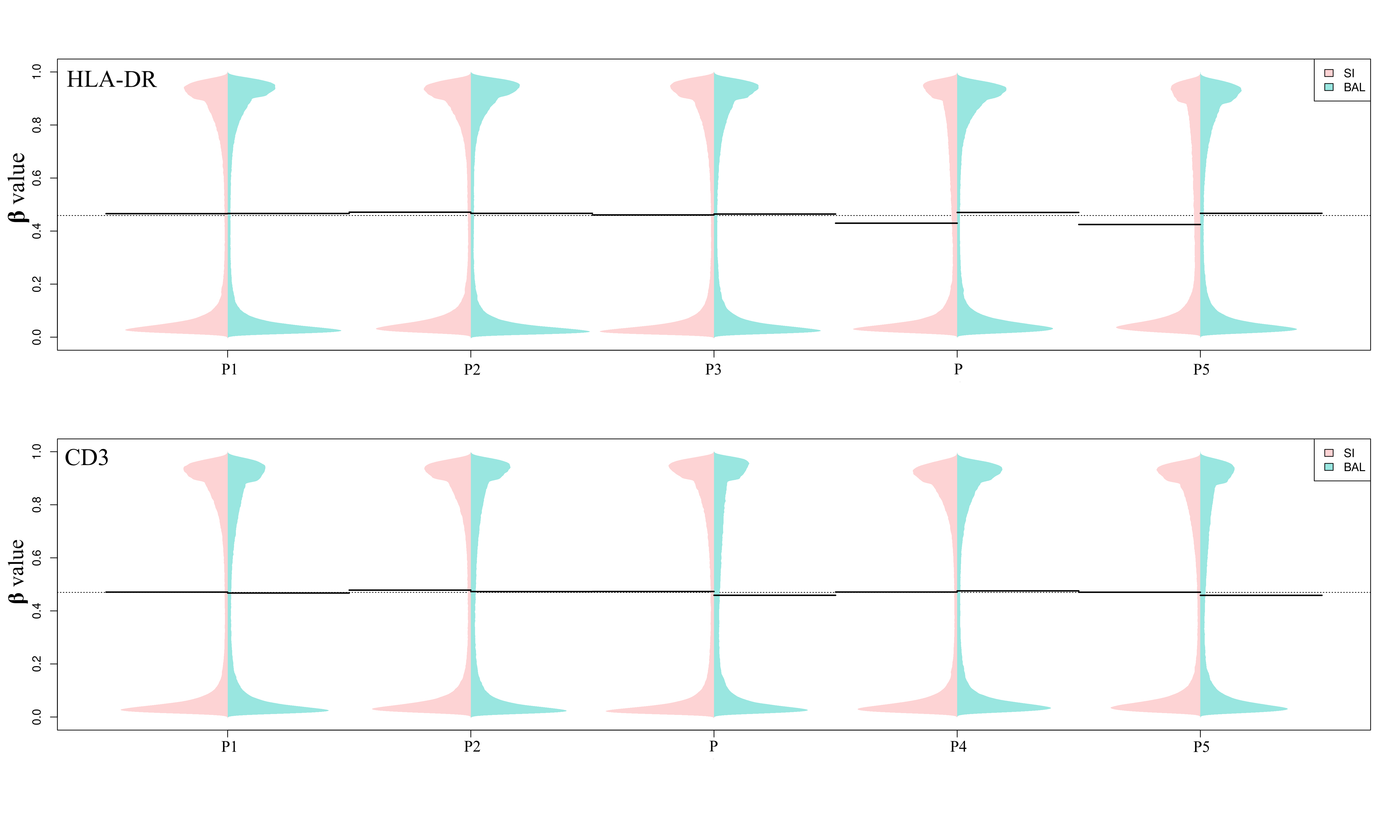

### Supplementary Figure S1b

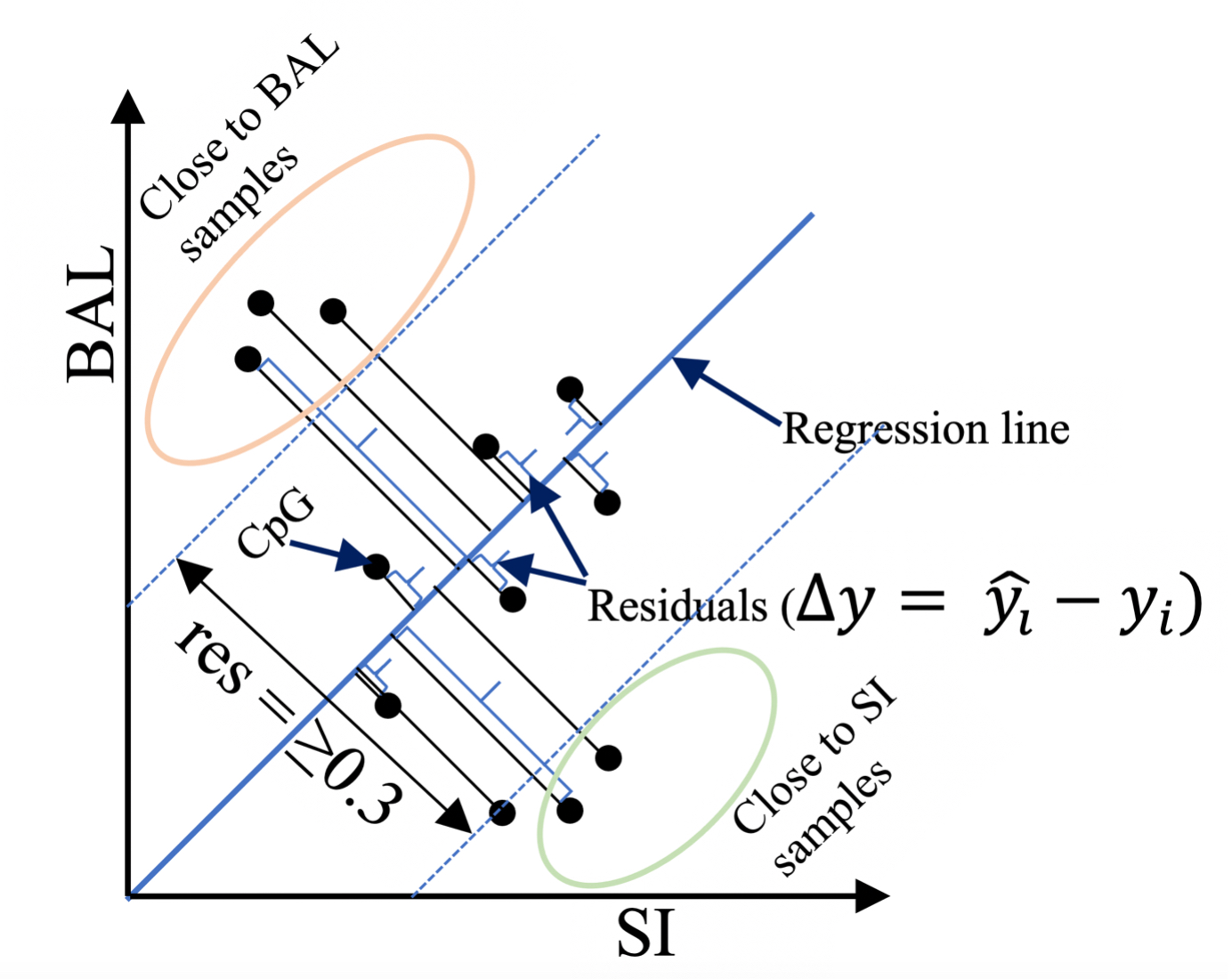

### Supplementary Figure S2

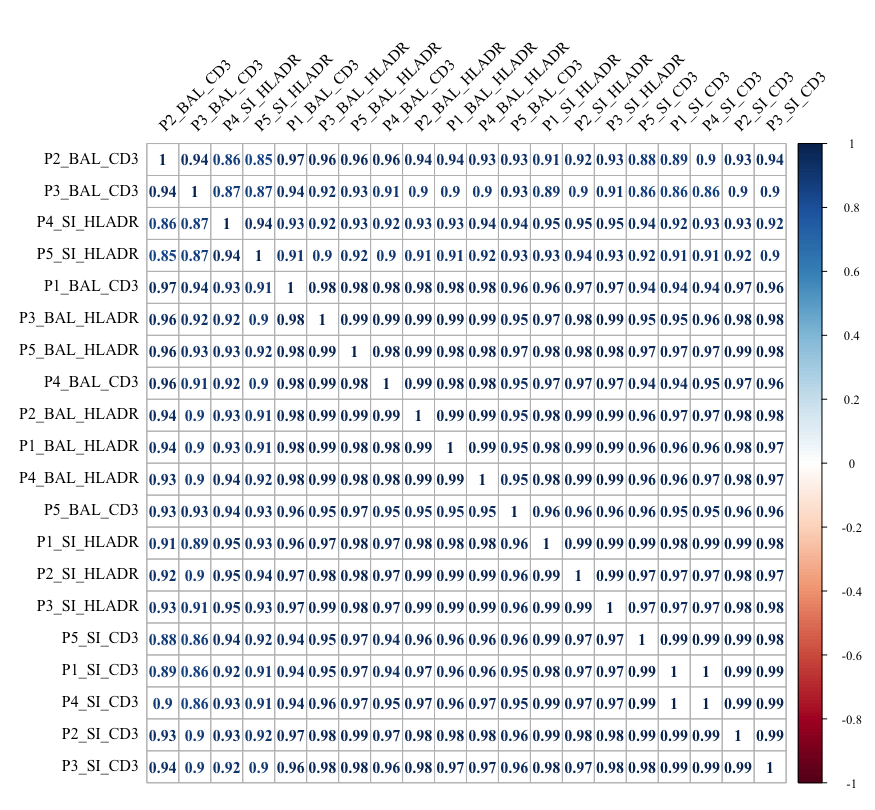
